# Physical, chemical, and structural properties of human gastric organoid-derived mucus

**DOI:** 10.1101/2025.11.08.687257

**Authors:** Katrina N. Lyon, Barkan Sidar, Cameron Dudiak, Chloe Vermulm, Grace Jordan, Thomas A. Sebrell, Alexis Burcham, Luke Domanico, Richard F. Helm, Wentian Liao, Bret Davis, Jennifer Brown, Scott G. McCalla, James N Wilking, Rama Bansil, Diane Bimczok

## Abstract

The gastric mucus layer protects the epithelium from gastric acid and ingested pathogens. However, studies of human gastric mucus have been limited due to poor accessibility of native human mucus and the abundance of contaminants in these samples. Here, we explored the potential of human gastric organoids as models for mucus production. Immunofluorescence staining confirmed that the organoids produced mucus containing MUC5AC and MUC6. The luminal mucus had viscoelastic properties similar to those of native human gastric mucus, as determined by particle tracking microrheology. To collect organoid-produced gastric mucus, termed bioengineered gastric mucus (BGM), organoids were cultured as monolayers at the air-liquid interface (ALI), and apically-secreted mucus was harvested and analyzed by MUC5AC ELISA, proteomics, CryoFE-SEM, and bulk rheometry. BGM contained high-molecular weight molecules also found in native gastric mucus, including MUC5AC. Proteomic analysis confirmed that BGM contained MUC5AC, MUC6, MUC1, and other stomach-specific molecules such as gastricsin, olfactomedin 4, and gastrokine. CryoFE-SEM showed that both BGM and native mucus had a porous structure and a characteristic honeycomb scaffold. Bulk rheometry confirmed that BGM exhibited shear thinning and predominantly elastic behavior, consistent with native mucus. Collectively, these findings indicate that BGM is an accessible alternative to native gastric mucus that can be produced on-demand for *in vitro* studies.

**New and Noteworthy:** We demonstrate the structural and functional similarities of organoid-derived gastric mucus and native mucus collected from human patients. The bioengineered gastric mucus mimics its native counterpart in its proteomic profile, physical architecture, and viscoelasticity. This work highlights the translational potential of organoid-derived mucus for functional investigations of the human gastric mucus layer.

## Introduction

The gastric mucus layer plays a crucial role in protecting the gastric epithelium from the harmful effects of orally-acquired pathogens such as *Helicobacter pylori* and from gastric acid (1). Mucus is a viscoelastic gel composed of large, hydrophilic mucin glycoproteins combined with electrolytes, lipids, and other smaller proteins (2). Mucin glycoproteins are the functional building blocks of mucus, with their capacity to bind large amounts of water to form hydrogels. Gastric mucus is composed of two major secreted mucins, MUC5AC and MUC6, which are produced by mucus pit cells and mucus neck cells, respectively, and one membrane-associated mucin, MUC1 (3). The composition, internal architecture and viscoelastic properties of the gastric mucus are considered essential for its function as a protective barrier (4, 5).

Gastric mucus has been difficult to study due to the limited accessibility of healthy human mucus samples and the inherent heterogeneity of fresh human mucus. Native, unpurified gastric mucus collected from animal or human stomach is commonly contaminated with microbiota (6), cell debris (7), and environmental contaminants (8), and purification procedures affect mucus structure and other properties (9). Moreover, native gastrointestinal mucus is subject to degradation by digestive enzymes such as pepsin and bacterial proteases, rendering it unstable (10, 11). Therefore, our current knowledge of gastric mucus is largely based on studies with highly processed porcine gastric mucin, which can be isolated and purified in large quantities from slaughterhouse material and which is commercially available (12). Studies with porcine mucin have led to key discoveries such as the ability of *H. pylori* to adhere to gastric mucus, to sense its mechanical properties, and to utilize urease to de-gel mucus through local pH changes (13–16). However, purified porcine mucin does not accurately replicate the chemical composition and structural complexity of the human gastric mucus layer *in situ* (9, 17). Specifically, processing and purification of mucin from native pig stomach disrupts disulfide bonds (18), which significantly alters the gel forming ability of the mucus and its biomechanical properties. Therefore, there is a need for improved models of the gastric mucus layer that replicate the native composition and structure of the mucus and that are also clean and sterile for experimental use (19–21).

Human organoids have been widely used as *in vitro* models of gastrointestinal homeostasis and disease, including *H. pylori* infection (22, 23). In a previous study, Sebrell *et al.* reported the presence of an optically dense substance within the organoid lumen as observed via backscatter light imaging, representing accumulation of mucus (24). Other studies from multiple groups have shown the presence of MUC5AC-positive pit cells and MUC6-positive neck cells in human and murine stomach organoids using immunohistochemistry, flow cytometry, and RNASeq (25–28). More recently, Boccellato *et al.* demonstrated robust mucus production by human gastric organoids cultured as monolayers at the air-liquid interface (ALI), which they termed “mucosoids” (20, 29). Similarly, the Allbritton group used murine organoid-derived ALI cultures to recreate the colonic mucus layer *in vitro* and demonstrated that the organoid model replicated key physiological features of native mucus (30, 31). Taken together, these data indicate that organoids may be a suitable source of sterile gastric mucus that does not require additional purification or processing steps that would disturb its structure and functionality.

To explore the potential of organoid-derived mucus for use in functional experiments, we here characterized the production, composition, and biophysical properties of mucus produced by 3-D human gastric organoids and organoid-derived monolayers cultured at an air-liquid interface (ALI). Our work provides a foundation for future studies with organoid-derived, bioengineered mucus (BGM) as a model to investigate physiological functions of mucus, cytoprotective treatments, and *H. pylori* infection.

## Results

### Human gastric organoids secrete mucus into the lumen

Previous studies have shown that human gastric organoids derived from adult stomach tissue express the two major secreted gastric mucins, MUC5AC and MUC6 (25, 27, 32). To confirm that gastric organoids cultured in our laboratory secrete gastric mucus, we performed Alcian blue staining, which indicates the presence of acidic mucin glycoproteins (33). In formalin-fixed paraffin-embedded sections, accumulation of mucin was detected both on the surface of gastric body tissue and in the lumen of the organoids (**Fig. 1A**). We next performed immunofluorescence staining for MUC5AC and MUC6. As expected, MUC5AC expression in human gastric body was restricted to the surface epithelium and gastric pit regions, consistent with MUC5AC expression by pit and surface mucus cells (**Fig. 1B**), whereas MUC6 expression was confined to the neck regions of the gastric glands (**Fig. 1C**). In gastric organoids, strong accumulation of MUC5AC was observed in the lumen. In contrast, MUC6 production appeared to be lower and mostly confined to the epithelial cell surface, with little accumulation. Overall, our findings confirm previous reports that gastric organoids produce mucus.

**Figure 1.**
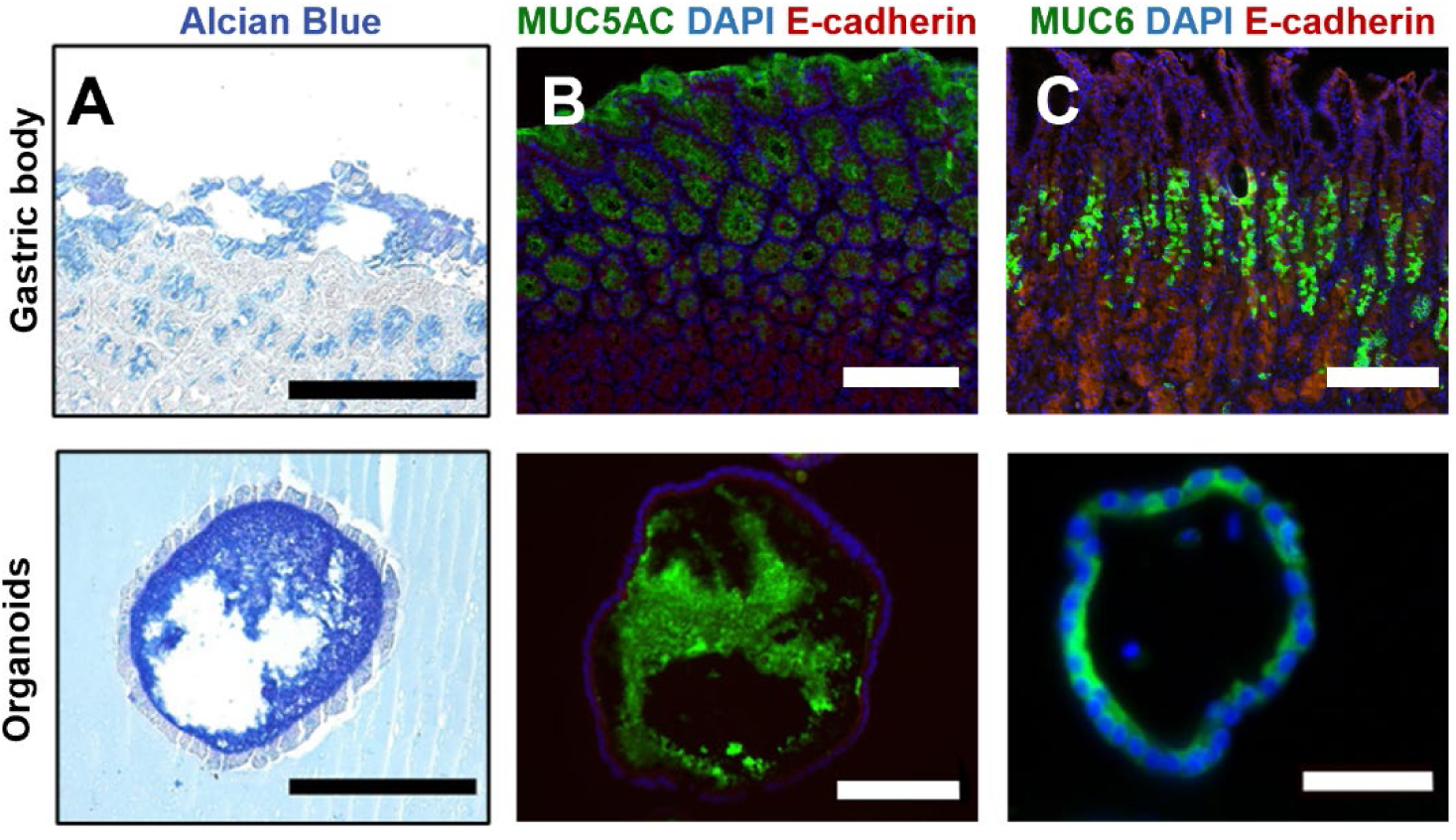
Polarized gastric organoids secrete and accumulate apical mucus. (**A**) Alcian blue staining for acidic mucins on tissue surface epithelium (top) and in the organoid lumen (bottom). Representative of 3 experiments. (**B**) Immunofluorescence staining for MUC5AC (green) on the surface epithelium and in the pit regions of human gastric tissue (top) and in the organoid lumen (bottom). Representative of two experiments. (**C**) Immunofluorescence staining for MUC6 in the neck regions of the gastric glands and on the outer edges of the organoid lumen. Representative of 2 experiments. All scale bars = 200 μm.

### Mucus that accumulates in 3-D gastric organoids is viscoelastic

We used live confocal imaging with subsequent particle tracking microrheology (34) to interrogate the viscoelastic properties of the luminal mucus in the gastric organoids. The organoids are topologically closed, embedded in Matrigel, and no significant active motion occurs in the lumen; thus, advection is limited and particle motion in the lumen is driven primarily by thermal energy (35). We injected red or green fluorescently-labeled colloidal polystyrene particles (d = 0.5, 1 µm) into multiple organoids, incubated them for 24 h, and then imaged the fluorescent particles for 30 sec at 26 frames per second with confocal time-lapse videomicroscopy at 24, 48 and 72 h (**Fig. 2A,B**). For each high frame-rate series, we used particle tracking image analysis to extract the 2D positional coordinates of individual particles over time and construct individual particle trajectories (**Fig. 2C**).

**Figure 2.**
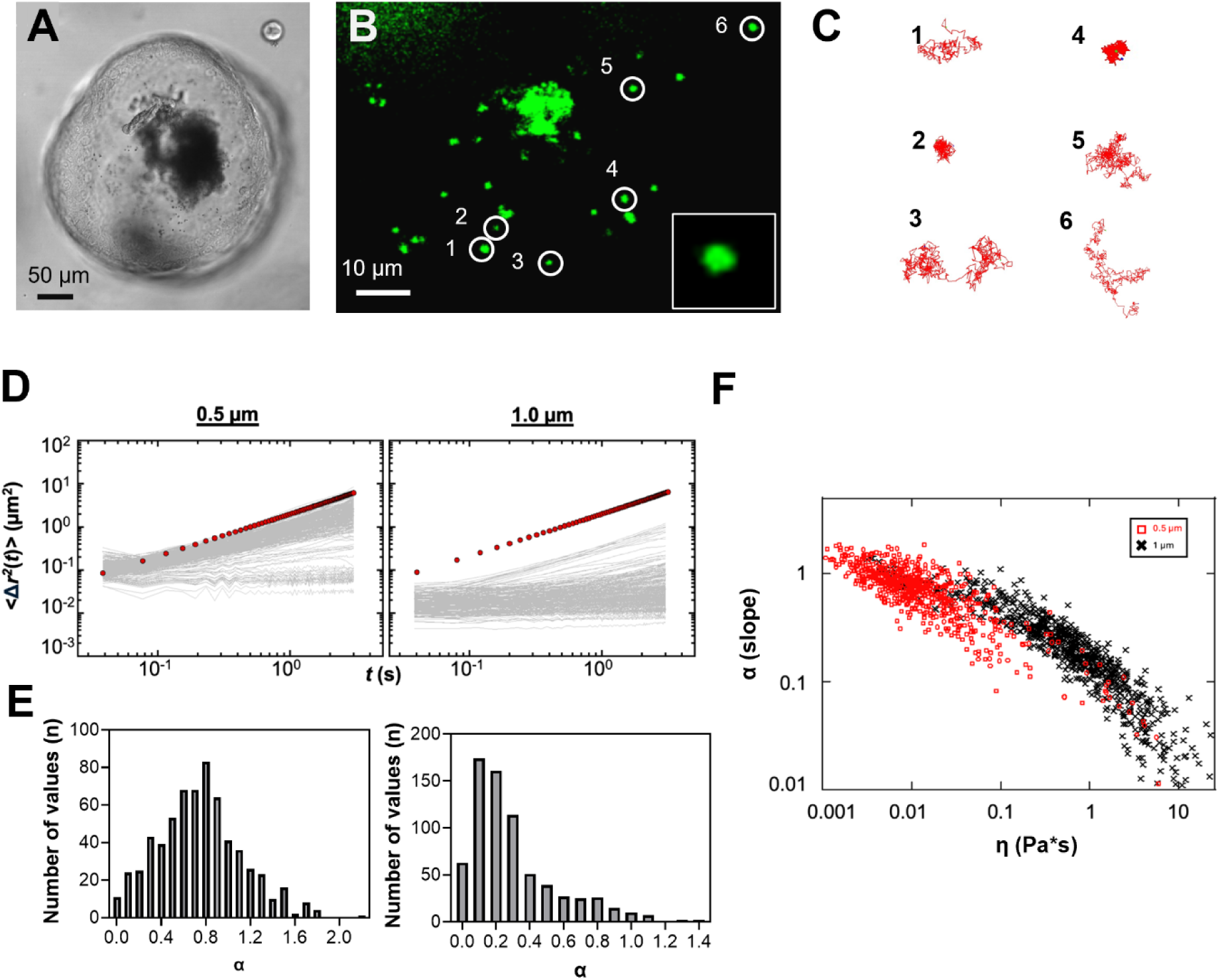
Particle tracking microrheology confirms the presence of a viscoelastic material in human gastric organoids. (**A**) Brightfield microscopy image of human gastric organoid microinjected with fluorescent polystyrene beads (d ≈ 1 µm). (**B**) High magnification confocal image of the organoid lumen from (A) shows multiple green fluorescent particles (d = 1.0 µm). White arrows indicate single particles. Inset: High resolution image of single particle. (**C**) Representative particle trajectories extracted from 30 sec videos captured on a confocal laser scanning microscope. Trajectories correspond to particles indicated with arrows in (B). (**D**) Time-dependent mean-squared displacement for individual polystyrene beads (left: d=0.5 µm, right: d=1.0 µm) that were imaged in one representative organoid out of nine organoids 24 h after injection. Red circles correspond to expected MSD for bead motility in water and were included as a reference (α = 1). (**E**) Motility distribution α for 0.5 µm (left, n=645) and 1.0 µm particles (right, n=717), representing the log slope of the mean squared displacement, of beads captured in five representative organoids each at 24 h, 48, and 72 h. (**F**) Relationship of α as a function of mucus viscosity (η, Pa*s). Red symbols represent *d =* 0.5 µm particles, and black symbols represent *d = 1* µm particles.

We found that particle diffusion was highly varied within a single organoid. The diffusion of some particles was confined, while others moved more freely, as illustrated by the trajectories shown in **Fig. 2C**. This variability indicates that the luminal environment in the organoids was spatially heterogenous. To quantify particle diffusion (or motility), we calculated the time-dependent mean-squared displacement (MSD), <Δr^2^(t)> of each particle and plotted these data as a function of time (**Fig. 2D**)(36). For a freely diffusing particle in a low viscosity fluid (i.e., water, indicated with red circles), <Δr^2^(t)> should scale as t^α^, with the diffusion exponent α = 1. Sub-diffusive scaling (α < 1) indicates that particle movement is constrained by the local microenvironment, enabling microrheological analyses. We determined the time dependent scaling α for 0.5 µm and 1 µm particles in five representative organoids (**Fig. 2E**). The resulting histogram distribution for 0.5 µm particles with a median α = 0.74 and a mode α = 0.8 indicated that most of these smaller particles were moving freely in the mucus (37). By contrast, α values for the 1 µm particles were significantly lower (*P* ≤0.0001; Student’s *t* test), with a median α = 0.22 and a mode α = 0.1, indicating that most 1.0 µm particles were confined in the mucus. Further, we plotted the viscosity (η, Pa*s) of the mucus as a function of α to find a negative slope (**Fig. 2F**). We observed that at a higher viscosity, the exponent α was lower, consistent with the expectation that the particles will move less in the higher viscosity regions.

Particles of 1 µm were used for subsequent analyses, as a majority of the 0.5 µm particles were moving freely, yielding less robust results. We next determined the storage (G’) and loss (G”) moduli of the luminal mucus at 1 Hz to assess viscous versus elastic behavior (**Fig. 3A, B**). Representative frequency-dependent storage and loss moduli are shown in **Suppl. Fig. 1**. For all mucus samples and time points, the median values for G” / G’ (tan δ), representing the ratio between the viscous and elastic moduli, were between < 1, indicating predominantly elastic behavior of the mucus, with a significant decrease in tan δ at 72 h compared to 48 h (**Fig. 3C**). Calculated median viscosity values were between 0.2 and 1, with a significant increase between the 48 h and 72 h time points (**Fig. 3D**).

**Figure 3.**
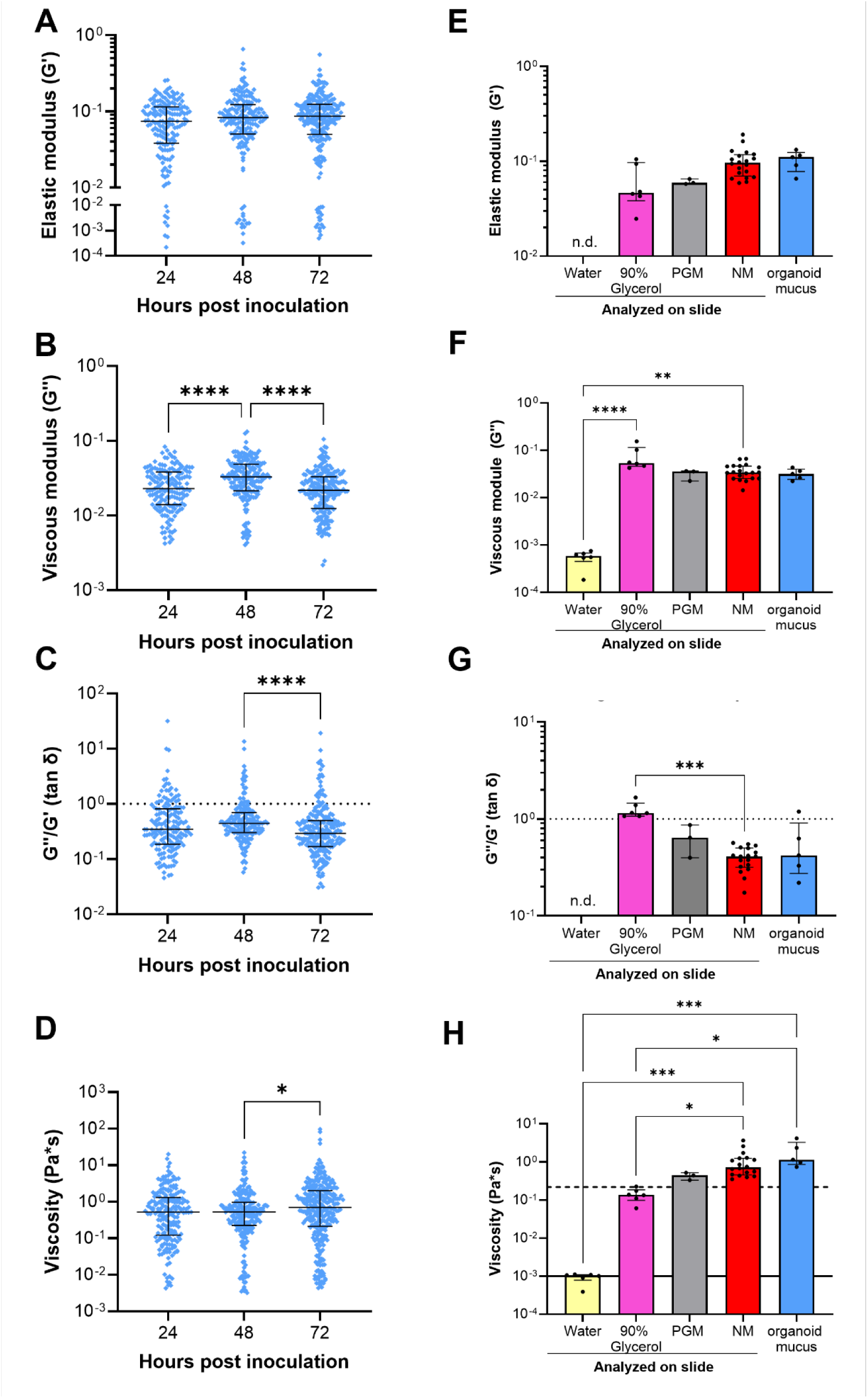
Rheological properties of mucus in gastric organoids are similar to those of native gastric mucus and are stable over time. (**A,B**) Elastic (G’) and viscous (G”) moduli calculated from the imaged particle displacement (low shear, 1 Hz) from five representative organoids at 24 h, 48 h and 72 h post injection. Individual data points, median, and interquartile range are shown. (**C**) Ratio of G’’/G’ (tan δ), where >1: viscous and <1: elastic. Kruskal-Wallis test with Dunn’s multiple comparison test; **** *P*≤0.0001. The dotted line represents the cutoff between predominantly viscous (tan δ > 1) and elastic (tan δ < 1) behavior. (**D**) Calculated low-shear (1 Hz) viscosity expressed as Pa*s. Kruskal-Wallis test with Dunn’s multiple comparison test; * *P*≤0.05. The X axis is set to the viscosity of water (10^-3^ Pa*s)(38). (**E, F**) Elastic (G’) and viscous (G”) moduli calculated from the imaged particle displacement (low shear, 1 Hz). Moduli are also shown for 90 % glycerol and 15 mg/mL PGM. (**G**) Tan δ values for water, 90% glycerol, 15 mg/mL PGM, and native mucus. Dashed line corresponds to cutoff between viscous (top) and elastic (bottom) behavior. (**H**) measured viscosity for water (reference value: solid line), 90% glycerol (reference value: dashed line), 15 mg/mL PGM, and native mucus. Data are from 3-6 independent videos each; three native mucus samples were analyzed and pooled data are shown. Organoid mucus shows averages for all time points for the five organoids represented in **A-D**.

For comparison, we analyzed porcine gastric mucin (PGM) and native human gastric mucus (NM) collected from discarded surgical tissues. Particle tracking microrheology was performed as described above with mucus samples in sealed microscope imaging chambers (**Fig. 3E-H**). Water and 90% glycerol were used for method validation and showed values in the expected range (38, 39). Importantly, we found that the viscous and elastic moduli, viscosity, and loss characteristics (tan δ) of the mucus in 3-D organoids were similar to those of PGM and native human gastric mucus (**Fig. 3E-H**). Taken together, these results indicate that the mucus in the human gastric organoids resembles native gastric mucus, exhibiting a heterogeneous viscoelastic behavior, with predominantly elastic characteristics.

### Production of bioengineered gastric mucus (BGM) on organoid-derived gastric epithelial monolayers cultured at the air-liquid interface (ALI)

To collect organoid-derived gastric mucus for functional experiments, gastric organoids were expanded, disaggregated, and reseeded on transwell inserts to generate organoid-derived epithelial monolayers (21) (**Fig. 4A**). The monolayers were then maintained in submerged culture until they reached confluence and developed a transepithelial electrical resistance of >200 Ωcm^2^ (TEER, **Fig. 4B**). To promote mucus secretion and accumulation, apical media was then removed to initiate an air-liquid interface (ALI, **Fig. 4C**), as described by Wang *et al*.(21). Apical mucus–which we term bioengineered gastric mucus (BGM)–was collected every 2-7 days (**Fig. 4D, E**), and volume and dry weight were measured. On average, monolayers produced 50±27 µL of BGM per well every 2 days (**Fig. 4E**). The dry weight of the BGM was 38.3 ± 22.0 mg/mL, considerably lower than the dry weight of native mucus (84.3 ± 58.1 mg/mL; **Fig. 4F**), but was in the expected range for an approximate water content of 95% for gastric mucus (40). Using an ELISA, we show that the BGM was rich in MUC5AC, the major mucin present in gastric mucus. MUC5AC concentrations in BGM samples were highly variable (0.265±0.429 ng/mL) but were in the same range as those in native gastric mucus collected from surgical tissue samples (NM, 299±0.497 ng/mL) (**Fig. 4G**). Next, size exclusion chromatography with multi-angle light scattering (SEC-MALS) was used to determine the molecular weight distribution of the BGM compared to NM. We found that molecular weight distributions of BGM and NM also were not statistically different, both samples showing high molecular weight compounds around 1.1×10^7^ g/mol for BGM and 1.6×10^6^ g/mol for NM and low molecular weight components around 8.2×10^4^ g/mol for BGM and 4.4×10^4^ g/mol for NM (**Fig. 4H, I**). Porcine gastric mucin (PGM) contained similarly sized high molecular weight compounds (3.6×10^6^ g/mol) as the BGM and NM, but lacked low molecular weight compounds (**Fig. 4I**). Overall, these data confirm that human gastric organoids, when cultured as monolayers at the ALI, can produce a MUC5AC-rich mucus layer that can be continuously harvested for functional analyses and has enhanced complexity compared to PGM.

**Figure 4.**
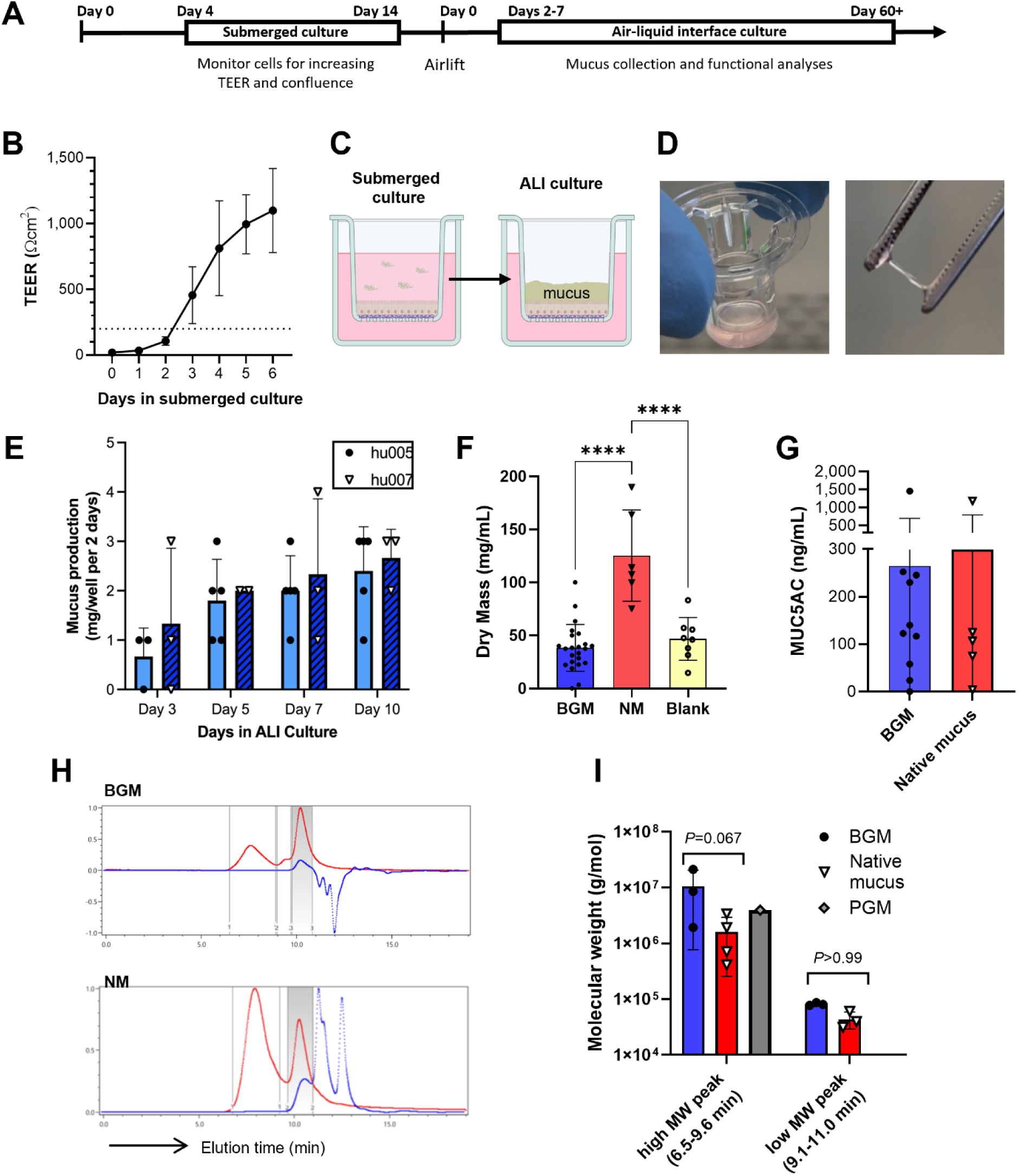
Production of bioengineered gastric mucus (BGM) on organoid-derived gastric epithelial monolayers cultured at an air-liquid interphase (ALI). **(A**) Timeline for 2D culture conditions, mucus collection, and functional analyses. (**B**) Time-dependent development of transepithelial electrical resistance (TEER); one representative out of seven experiments with four technical replicates. Mean ± SD; dotted line indicates TEER threshold for airlift at y=200 Ωcm^2^. (**C**) Schematic diagram of Transwell 2D culture conditions: submerged (left) and air-liquid interface, ALI (right). (**D**) Clear, viscoelastic mucus on the Transwell insert (left) and removed from the apical epithelium (right). (**E**) Bioengineered mucus (BGM) dry mass collected per well every other day (mean ± SD), representative of two experiments with 3-5 technical replicates. Dry mass was determined by weighing lyophilized samples. (**F**) Concentration (mg dry weight per mL) of BGM (n=23), native mucus (n=6), and L-WRN culture media (n=8). Data analyzed by one-way ANOVA with Tukey’s multiple comparisons test; \*\*\*\**P* ≤ 0.0001. (**G**) Concentration of MUC5AC in BGM (n=10) and NM (n=5) determined by ELISA. (**H**) Size exclusion chromatography paired with multi-angle light-scattering (SEC-MALS) analysis of BGM (top) and NM (bottom) showing the presence of low and high molecular weight compounds based on differential refractive indices (dRI) in blue and the light scattering (LS) signal in red. (**I**) Molecular weights of a porcine gastric mucin (PGM) control, BGM (n=3), and NM (n=4) determined by SEC-MALS. Data analyzed by one-way ANOVA with Tukey’s multiple comparisons test. All bar graphs include individual data points, mean ± SD.

### Proteomic analysis of bioengineered gastric mucus reveals physiologically relevant secretome

We next determined the biochemical composition of the BGM samples (n=3) and compared them to native mucus (NM, n=4) and PGM (n=1). Using mass spectrometry, we identified between 161 and 196 proteins per sample in BGM, and between 216 and 1,611 proteins in NM (**Fig. 5A**). Across all samples, a total of 2,055 unique proteins were identified in native human gastric mucus, but only 58 were present in samples from all four donors. In contrast, only 304 unique proteins were found in BGM (n=3), but 74 were present in all three samples. Overall, 130 proteins were detected in both BGM and NM samples (**Fig. 5B**, top). Only 75 proteins were found in PGM (n=1), sharing 27 with the BGM samples (**Fig. 5B**, bottom).

**Figure 5.**
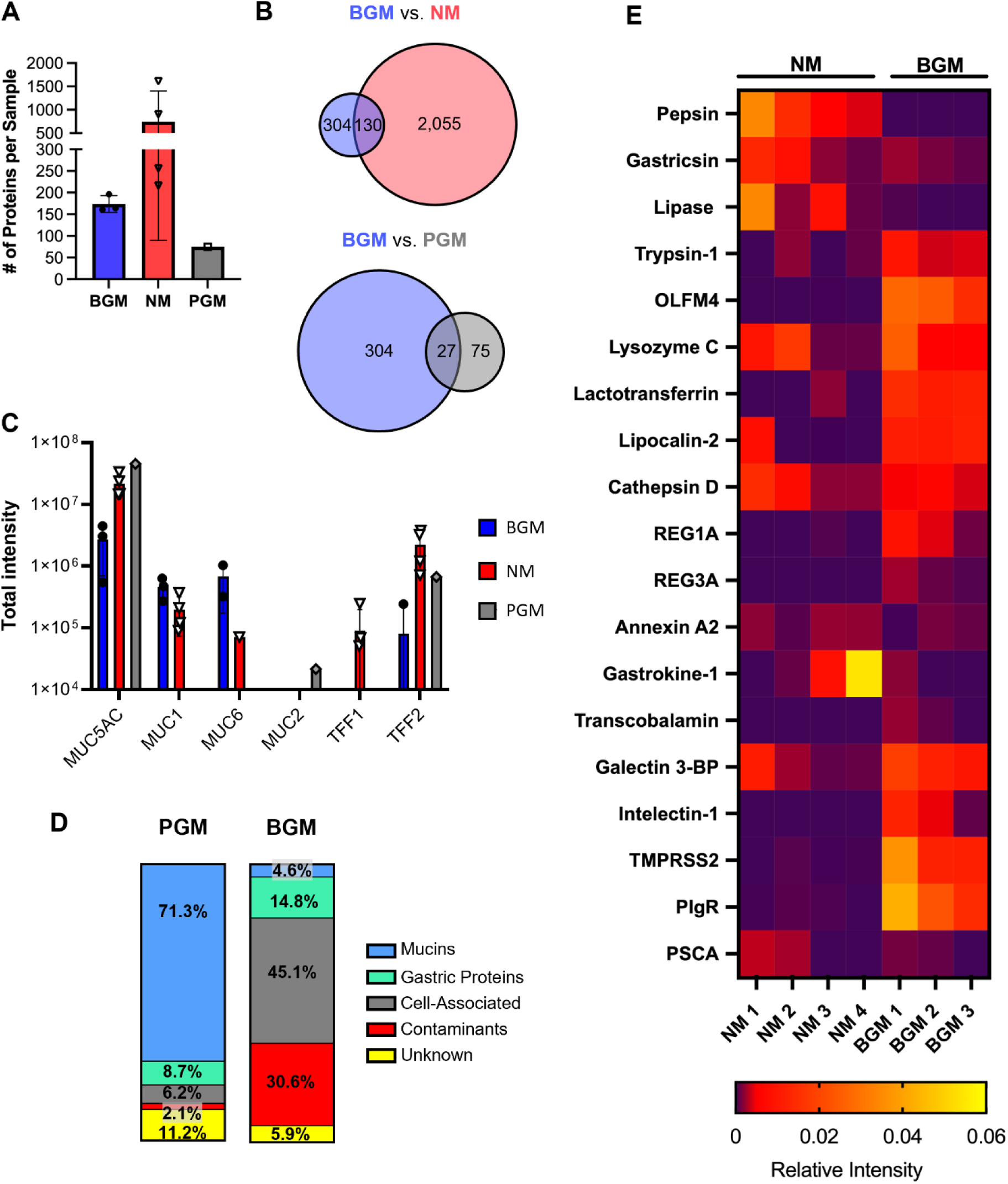
Proteomic analysis of bioengineered gastric mucus. Three biological replicates of BGM, four native human gastric mucus samples collected from surgical material, and one porcine gastric mucin sample were analyzed by LC/MC. (**A**) Number of proteins identified by mass spectrometry in bioengineered gastric mucus (BGM; n=3), native mucus (NM; n=4), and porcine gastric mucin (PGM; n=1) samples. (**B**) Number of total and shared proteins (overlapping region) between BGM and NM (top) and BGM and PGM (bottom). (**C**) Relative abundance of gastrointestinal mucins MUC5AC, 1, 6, and 2 and trefoil factors (TFF) 1 and 2 in BGM (blue), NM (red), and PGM (grey). Combined data from all identified protein isotypes. (**D**) Categorization of proteins detected in PGM (left) and BGM (right) based on protein functions and cellular distribution listed in the Human Protein Atlas (79). Percentage values represent cumulative relative protein intensity values for each category. (**E**) Heatmap showing relative expression of key stomach-specific factors identified in BGM and NM. Data from n=4 NM and n=3 BGM samples.

MUC5AC, MUC1 and MUC6 all were among the top 35 identified proteins in BGM (**Fig. 5C, Supplementary data 1).** Comparison between the BGM and NM revealed a high variability in the abundance of MUC5AC, MUC1, or MUC6 within and between sample types (**Fig. 5C**). In PGM, only MUC5AC and the intestinal mucin MUC2 were detected, but not MUC6 or MUC1. Interestingly, trefoil factor 1 (TFF1) was only detected in NM, whereas TFF2 was present in NM, BGM, and PGM. The 35 most abundant proteins detected in all three BGM samples included epithelial-derived immunoactive substances such as lysozyme, complement factor C3, and lactotransferrin. In contrast, the top 35 most highly expressed proteins present in all NM samples included functionally important gastric proteins such as lipase, carbonic anhydrase, trefoil factors 1 and 2, and antibody fragments, which were absent from BGM. Both BGM and NM contained cytoskeletal proteins such as actin, keratins, and cytokeratins and cellular enzymes, indicative of cellular contamination. Even in the highly purified porcine gastric mucin, contaminants such as hemoglobin and cellular metabolic enzymes such as phosphopyruvate hydratase and aspartate aminotransferase were among the top 35 most abundant proteins that were identified (**Supplementary data 1**).

A comparison of protein groups between PGM and BGM (**Fig. 5D**) showed that, based on total intensity values, PGM consisted mostly of mucins (71.3%), with 8.7% other gastric proteins, and the remaining proteins were cellular proteins, contaminants, and unknown proteins. BGM only contained 4.6% of mucins, but a higher proportion of other gastric proteins than PGM, at 14.8%. However, BGM was significantly contaminated with cellular proteins (45.1%), other contaminants (30.6%) and unknown proteins (5.9%).

When focusing on stomach-specific proteins other than mucins, a comparison between BGM and NM based on relative intensity values found that BGM was lower in gastric enzymes pepsin, gastricsin, and lipase, but showed increased expression of other gastric proteins such as olfactomedin 4 (OLFM4) and immunoactive factors such as lipocalin 2, lactotransferrin, poly-immunoglobulin receptor (pIgR), and galectin 3 binding protein (galectin 3-BP). Overall, these data show that BGM contains major gastric mucins and other functional gastric proteins, including several factors that are not found in PGM, but that it differs from both native mucus and porcine mucin in protein composition and abundance and presence of cell and tissue contaminants.

Visualization of bioengineered mucus by cryo-field emission scanning electron microscopy shows a characteristic porous architecture.

To assess the structure of BGM and to compare BGM to native human gastric mucus, we used cryo-field emission scanning electron microscopy (CryoFE-SEM) followed by image analysis using a Segment Anything Model (41). As shown in **Fig. 6A**, both native mucus and BGM had similar, smooth surface patterns that were continuous aside from the fracture lines. Fracturing of the samples immediately prior to imaging exposed visually similar cross-sections with honeycomb-like scaffolds and pores in both BGM and NM (**Fig. 6A, Suppl. Fig. 2A**). After correction for outliers, digital image analyses revealed that honeycomb cells were significantly larger in BGM, with a median area of 38.8 µm^2^ compared to NM, where honeycomb cells had a median area of 32.0 µm^2^ (**Fig. 6B**). Pores were also significantly larger in BGM, with a median area of 0.86 µm^2^, compared to NM, where the median pore area was 0.43 µm^2^ (**Fig. 6C**). To confirm that the microscopic structures observed were not also present in tissue culture media, we imaged L-WRN cell culture medium alone and L-WRN medium collected from blank, collagen-coated Transwell® inserts (42) (**Supp. Fig. 2B**). These controls had morphologies clearly distinct from the mucus samples in that they lacked both rounded pore structures and the pseudohexagonal geometries of the honeycomb scaffolds seen in BGM and NM. These data confirm the internal architecture of our BGM to be consistent with NM as well as other types of mucus when visualized with CryoFE-SEM (43–45).

**Figure 6.**
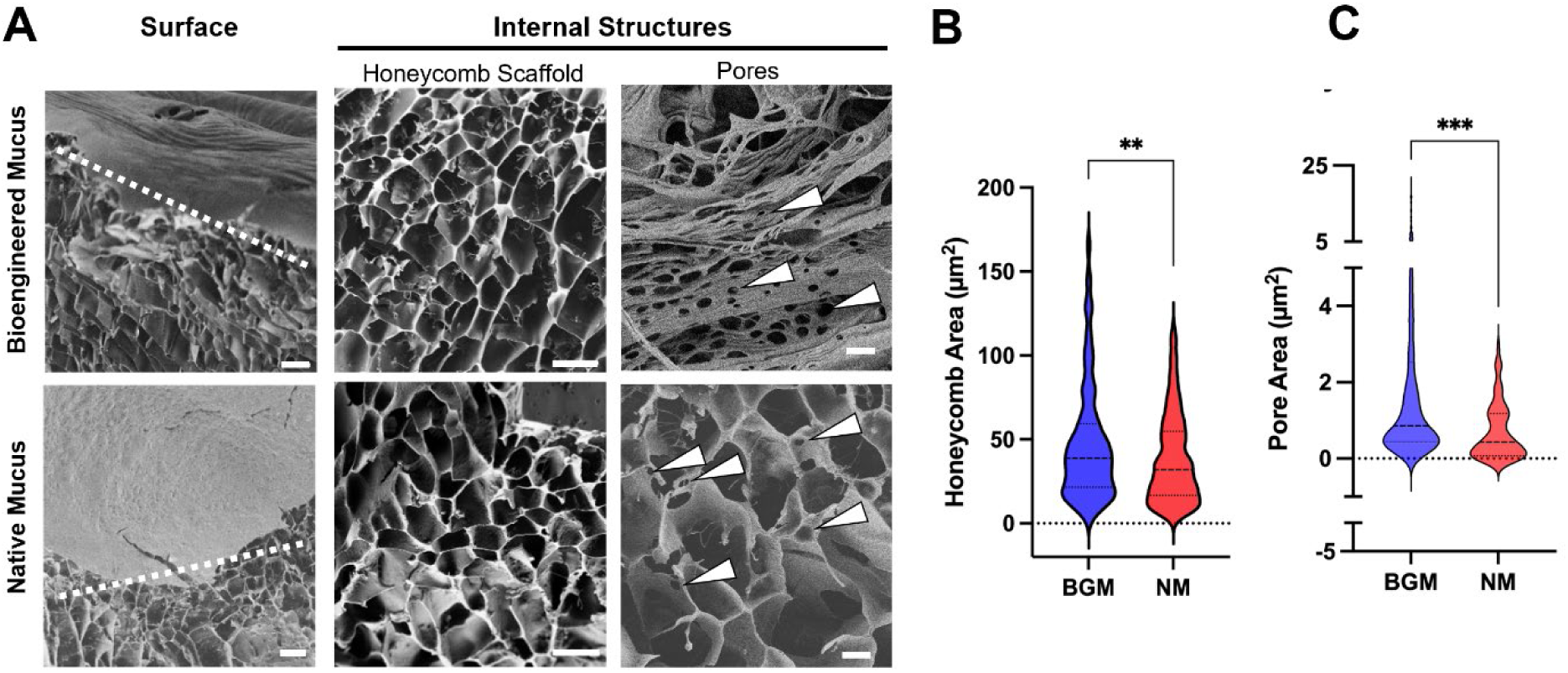
Visualization of bioengineered gastric mucus by cryo-scanning electron microscopy. (**A**) Left: Electron micrographs showing frozen surfaces of BGM (top left) and NM (bottom left) with fractures (roughly indicated by dotted lines) to expose internal architecture (scale bar = 20 µm). Middle: Electron micrographs of internal honeycomb scaffold in BGM (top middle) and NM (bottom middle) (scale bar = 10 µm). Right: Pores in BGM (top right; scale bar = 5 µm) and NM (bottom right; scale bar = 5 µm). Arrows point to representative pores. (**B**) Honeycomb measurements for BGM (n=166 cells from 2 biological replicates) and for NM (n=362 cells from 2 biological replicates). Student’s t test: ** *P =* 0.0018. (**C**) Pore mesh area for BGM (n=380 pores from 3 biological replicates) and NM (n=60 pores from 2 biological replicates). Student’s t test: *** *P =* 0.0004. Pores and honeycomb areas were measured by ImageJ using SAM, and all outliers were removed using the ROUT coefficient with a Q value of 1%.

### Rheometric frequency sweeps confirm viscoelastic behavior of bioengineered mucus

For rheological characterization of the BMG samples, we measured the steady shear viscosity and frequency-dependent loss and storage moduli of BGM and NM samples using an AR-G2 rheometer with a 20 mm parallel plate geometry (**Fig. 7A,B**). Flow sweeps between 1 and 100 Hz to assess overall viscosity showed consistently higher viscosity for NM than for BGM (**Fig. 7A**). The BGM and NM had similar trends in their frequency-dependent viscous (G’’) and elastic moduli (G’) (**Fig. 7B**). The slopes of the elastic moduli for NM and BGM, respectively, were 0.3285 (*p*=0.0008) and 0.4353 (*p*=0.0040). The slopes of the viscous moduli, similarly, were 0.08349 (*p*=0.0072) and 0.09155 (*p*=0.0293). The values for tan δ—the ratio of viscous to elastic moduli—demonstrated that the NM and BGM both had predominantly elastic behavior (**Fig. 7C**).

**Figure 7.**
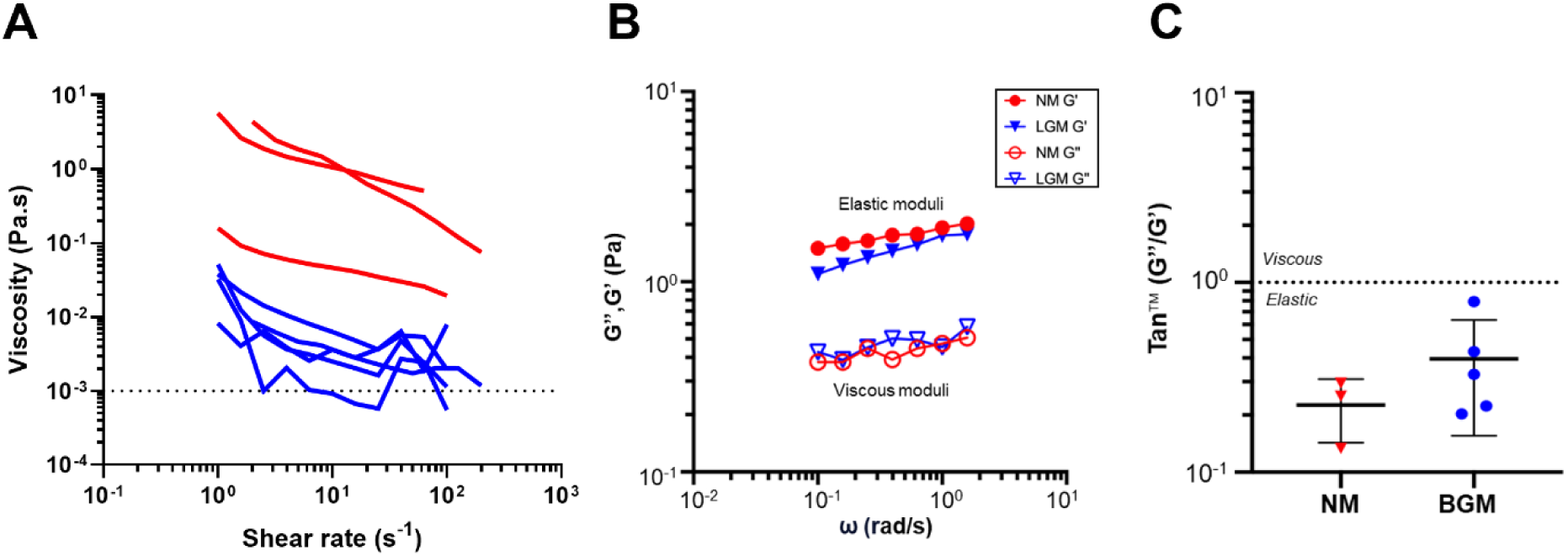
Rheological comparison of bioengineered and native gastric mucus. (A) Steady shear viscosity as a function of shear rate for bioengineered gastric mucus (BGM; blue; n=5) and native mucus (NM; red; n=3). Dotted line represents the viscosity of water (10^-3^ Pa·s) as reported by Lai *et al*.(53) (B) Frequency-dependent elastic (storage, G’) and viscous (loss, G’’) moduli of NM (red) and BGM (blue) (n=1). Representative of 3 independent experiments. (C) Ratios of viscous to elastic moduli of NM (red; n=3) and BGM (blue; n=5) (>1: predominantly viscous behavior, <1: predominantly elastic behavior). Each point represents the average of tan δ over 0.1-1.5 rad/s, for each independent experiment. All frequency sweep data were performed within the linear viscoelastic region.

## Discussion

In this study, we explored human gastric organoids and organoid-derived monolayers as models of the human gastric mucus layer. Specifically, we used microscopy and particle tracking microrheology to analyze gastric mucus in the lumen of 3-D organoids. Furthermore, we provide a detailed physical, chemical, and structural characterization of gastric mucus harvested from human gastric organoid-derived monolayers cultured at an ALI.

Using Alcian Blue and immunofluorescence staining for MUC5AC and MUC6, we confirmed the mucin content of the luminal material in the 3D organoids. We also showed that the mucus inside the 3D organoids had rheological characteristics that were similar to those of native gastric mucus (NM) and porcine gastric mucin (PGM). To harvest the organoid-derived mucus, termed bioengineered gastric mucus, BGM, for functional experiments, we used a transwell monolayer model. Our data confirm studies from other laboratories showing that robust mucus production by organoids can be induced when cells are cultured at an air-liquid interface (ALI) (20, 29, 30, 46). Gastric organoid-derived ALI monolayers produced up to 3 mg of BGM per well every other day, with an average dry weight of ∼40 mg/mL. This is roughly equivalent to a 4% dry matter content, which is in the expected range (1).

Based on our results, we propose that BGM is a more physiological model of the human gastric mucus layer than the xenobiotic porcine gastric mucin (PGM) that is commonly used in experiments. Previous studies PGM have led to significant insights into gastric mucus function and its interactions with *H. pylori*, such as the ability of the mucus to undergo pH-dependent changes, and the ability of *H. pylori* to de-gel mucus via urease activity (14, 47, 48). However, purification of PGM involved numerous processing steps such as filtration and chromatography methods that result in the removal of potentially relevant compounds (18). Our proteomics analysis showed that PGM shared key mucus compounds with NM and BGM, including MUC5AC and TFF2. However, PGM lacked MUC6 and MUC1, two mucins produced by gastric epithelial cells that were present in NM and BGM (49, 50). We also found that PGM lacked small molecule compounds that were present in both NM and BGM, but contained MUC2, a mucin produced by intestinal epithelial cells that is rarely expressed in the healthy stomach epithelium (51). Likely due to the extensive processing steps involved in the purification of PGM (18, 52), the number of proteins identified was much lower, and cell associated proteins and other contaminants that were present in BGM and NM were lost. It is unclear to what extent the lack of these factors in PGM impacts its ability to serve as a physiologically relevant model of the human gastric mucus layer (53).

Importantly, we show that BGM replicates key features of NM, with important mucin components MUC5AC, MUC1, MUC6 detected in both sample types. The concentration of MUC5AC in the BGM–measured using an ELISA assay–was highly variable, but not significantly different from that in NM, with average concentrations around 250 – 300 ng/mL. In both BGM and NM, the amount of MUC5AC was markedly higher than that of MUC6, in agreement with previous findings (54). Our SEC-MALS analysis of mucus glycopeptides detected high molecular weight (MW) compounds with sizes around 1×10^6^ – 1×10^7^ g/mol in all three sample types, BGM, NM and PGM, but only BGM and NM also contained low MW compounds around 4-8×10^4^ g/mol. Squire *et al*. found components with molecular weights of only 7.2×10^5^ g/mol for human gastric mucus glycopeptides from the antrum using the same analytical method, but samples had been treated with a protease, which likely resulted in smaller molecules (54). Tandem mass spectrometry analysis revealed similar levels of gastricsin, lysozyme C, cathepsin, and galectin 3-BP in BGM and NM. Visualization of the internal structural architecture of the BGM and NM using cryoFE-SEM showed a honeycomb scaffold and pore structures that were consistent with those reported in other mucus studies (44, 45, 55–59). Additionally, elastic and viscous moduli and the loss factor tan δ were similar in BGM and NM, indicating comparable rheological characteristics.

Both NM and BGM were highly variable in features including dry mass, MUC5AC concentration, molecular weight distribution and number of proteins per sample. Notably, we discovered some differences between BGM and NM that should be taken into consideration. First, dry mass of the BGM was significantly lower, and the number of proteins was reduced. Second, the viscosity of the BGM measured by bulk rheology also was lower than that of NM. Third, the BGM also had significantly larger honeycomb-like structural domains and pores than NM. This enlarged network structure in BGM may indicate lower crosslinking density, which aligns with the consistently lower viscosity compared to NM. We hypothesize that the reduced dry weight, viscosity, and honeycomb and pore sizes indicate that some BGM samples used in our analyses may have been diluted by media contamination from the basolateral transwell chamber. This hypothesis is supported by the presence of tissue culture media components such as hemoglobins or albumin from serum in the BGM samples.

Proteomic analysis revealed additional differences in mucus composition between BGM and NM. Concentrations of gastric enzymes such as pepsin and gastric lipase were much lower in BGM. However, these enzymes could easily be added exogenously for experimental purposes. Interestingly, BGM contained increased levels of immunologically-active substances including lactotransferrin, lipocalin-2, olfactomedin, intelectin-1, the regenerating gene (REG) family members REG Iα and REG 3α, and the polymeric immunoglobulin receptor (pIgR). It has previously been reported that the gastric mucus layer contains antimicrobial factors (60). Our data confirm data by Vllahu et al. that showed secretion of lactotransferrin and lipocalin by gastric organoid-derived ALI monolayer cultures (46). Importantly, secretion of these antimicrobial factors was significantly enhanced upon epithelial cell stimulation with pro-inflammatory cytokines (46). These findings suggest that BGM may be a useful model for studying aspects of gastric mucus related to antibacterial and innate immune functions.

To the best of our knowledge, our study is the first to utilize particle tracking microrheology to probe the luminal compartment of organoids. Importantly, our microrheological analysis found that the calculated overall viscosity and the viscoelastic properties (tan δ) of luminal mucus inside 3-D organoids were very similar to measurements obtained for native human gastric mucus samples from three patients at neutral pH and for PGM used at a standard concentration of 15 mg/mL (14, 61). Notably, most of the smaller 0.5 µm particles experienced mostly free or slightly hindered diffusion, whereas the larger 1 µm particles appeared to be mostly trapped in the mucus gel. The differential motility of 0.5 and 1 µm particles was likely caused by the porous structure of the hydrogel, as shown by our cryo-FE SEM analysis. In a hydrogel, water fills the space between macromolecules (62), allowing unrestricted movement of small particles within these spaces. Previous studies on native mucus harvested from canine and porcine stomach showed that, at the microscale level, gastric mucus has a honeycomb structure with cells sizes of about 4 µm in diameter (equivalent to an area of 12.6 µm^2^) (43, 44). Image analysis of honeycomb cells in cryo-FE SEM images of in that the cells in the honeycomb scaffold were in the range of 30-40 µm^2^. However, smaller, round pores detected on the walls of the scaffold were generally <1 µm^2^, suggesting that beads of 1 µm diameter could become wedged in these pores, limiting diffusive motility. Importantly, the viscosity values we obtained for both luminal organoid mucus and native mucus using microrheology were similar to previously reported values for human gastric mucus at a low shear rate (approx. 3 −9 Pa *s at 1.15 Hz)(63). Moreover, the particle tracking microrheology data were highly reproducible and are therefore helpful for comparative analyses in samples where bulk methods are not feasible and may provide more relevant insights into mucus properties when considering interactions with micro-scale pathogens or drug formulations.

We are aware that our study has several limitations. As seen in the results, there was a large variability in the composition and mucin concentration of individual BGM samples, pointing to a need for consistent quality control steps prior to using BGM in functional experiments. It is unclear whether this large variability was caused by genetic or epigenetic differences between tissue donors and organoid cultures or by other factors, and therefore needs to be investigated in more detail. A second limitation is that we did not compare mucus characteristics at different pH values but instead only assessed samples at a neutral pH. These analyses are important second step experiments, given the well-known impact of pH on mucus gelation (47, 61, 64), but were beyond the scope of the current study. Notably, mucus in the organoid lumen is approximately neutral at baseline, as we have shown (65), and all tissue samples that human mucus was recovered from were stored in excess volumes of neutral, buffered tissue culture media for 16-24 h prior to mucus harvesting, which resulted in a neutral pH for the native mucus samples used in our study. A third limitation was the methodology used to prepare mucus sample for MS proteomics, which involved standard trypsin digestion to generate peptides. Because trypsin digestion has a lower efficacy in highly glycosylated samples, this approach may have led to an underestimation of the mucin content (66).

Overall, the BGM model has strong potential to serve as a useful *in vitro* model of the human gastric mucus layer, especially when using a well-defined BGM biobank to analyze the impact of bacterial infection, drug exposure, or other interventions. However, quality control steps should be implemented to ensure a consistent mucus composition and concentration. This could include using assays to check mucin concentration, or monitoring the color of the collected BGM for signs of media leakage. In future studies, the cellular composition and differentiation of gastric organoids could be adjusted by altering their culture conditions (20, 29). For example, gastric organoid cultures may be driven toward MUC5AC-producing foveolar cells by supplementing the media with epidermal growth factor (EGF), or toward a MUC6-dominant phenotype by removing EGF and adding bone morphogenic protein (BMP)(29).

## Materials and Methods

### Human Gastric Tissue and Mucus Samples

Human gastric tissue samples for the generation of organoid cultures were obtained with informed consent and IRB approval from patients undergoing endoscopy and biopsy at the Bozeman Health Deaconess Hospital (protocol 2023-48-FCR). Alternatively, de-identified whole stomachs collected post-mortem from transplant donors or surgical discard material from sleeve gastrectomies were provided by the National Disease Research Interchange (NDRI, protocol DB062615-EX). Tissue samples from the NDRI were transported to Montana State University via overnight shipping in DMEM on ice and were used for both organoid establishment and mucus collection. To collect native mucus from the tissue samples, the mucus was gently scraped off the luminal tissue surface using a clean glass slide. To ensure that the least-contaminated and most representative portion of the native mucus sample was used, the clearest portion of the mucus was removed from the tube with forceps, leaving behind any contaminating blood or gastric juice. Mucus that showed major contamination with blood or other cellular debris was discarded. Representative mucus samples (n=4) had a pH of 6.5±0.5, likely due to prolonged storage in buffered media during shipping.

### 3D Organoid Culture

Human gastric organoid cultures were established and maintained as previously described (67–69). Briefly, gastric glands were isolated by collagenase digestion and seeded into Matrigel (Corning). Polymerized Matrigel was overlaid with L-WRN culture medium containing the supernatants of murine L cells that secrete the growth factors Wnt3a, noggin, and R-spondin 3(70). The composition of the L-WRN culture medium is listed in **Supplementary Table 1** (67). Organoid cultures were passaged weekly and were generally used between passages 2 and 15.

### Alcian blue and immunofluorescence staining

For histological analysis of organoids, culture medium was aspirated, and the Matrigel was overlaid with Histogel^TM^ (Epredia HG4000012) which was allowed to polymerize for 15 minutes at room temperature in a biosafety cabinet. A mini cell scraper (Biotium #22003) was then used to transfer the Histogel^TM^ into a cassette lined with filter paper. The cassette was submerged in 10% neutral buffered formalin (Richard Allen Scientific, Kalamazoo, MI, USA) overnight, and then transferred to 70% ethanol for at least 24 hours prior to paraffin embedding on a Sakura Tissue-Tek VIP1000 tissue processor/embedding center. Gastric tissue samples were also fixed in formalin and then embedded in paraffin. 5 µm sections were then prepared on a Leica 2035 rotary microtome. For staining, slides were deparaffinized with xylene and ethanol in a fume hood. Sections were stained with an Alcian blue-acetic acid solution (pH 2.5, Sigma) for 30 minutes and counterstained with nuclear fast red for 5 min. For detection of mucins MUC5AC and MUC6 by immunofluorescence, slides were then treated to heat-induced epitope retrieval using Vector Antigen Unmasking Solution in a rice cooker (71). Sections were blocked for 30 min with animal-free blocking reagent (Vector Laboratories) and incubated with primary antibodies to E-cadherin (Biolegend Cat. 324102), MUC5AC (Santa Cruz Biotech Cat. 398985), and MUC6 (Invitrogen Cat. PA5-50557) overnight at 4°C. Samples were washed with DPBS and secondary antibodies (goat anti-mouse IgG2a AlexaFluor 488, Cat. 1080-32; goat anti-mouse IgG1 AlexaFluor 488, Cat. 1070-30; and goat anti-rabbit IgG1 AlexaFluor 555, Cat. 4050-32, respectively, all from Southern Biotechnology) were added for 2 h at room temperature with gentle agitation. After a final wash step, samples were stained with DAPI and covered with Fluoroshield (Sigma) for mounting. Standard brightfield or fluorescence imaging was performed using a Keyence BZ-X810 microscope.

### Particle tracking microrheology

For particle tracking microrheology in organoids, gastric organoids cultured on 35 mm MatTek glass bottom plates were microinjected with fluorescent microspheres. A 2 µL glass capillary was backfilled with sterile mineral oil and loaded onto a micromanipulator-controlled Nanoject (Drummond 3-000-204). The capillary was then filled with a 4.55 × 10^6^ particle/mL solution of 1 µm Fluoresbrite polystyrene microspheres (Polysciences 17154-10) in PBS. 9.2 nL of the particle solution was injected into organoids that had a diameter of at least 300 µm (n=10). The organoids were then incubated at 37 °C, 5% CO_2_ for 24 hours prior to imaging to allow equilibration of the system. Organoids were imaged on a Leica SP5 confocal laser scanning microscope on a heated stage with an environmental control chamber (37 °C, 5% CO2; Life Imaging Services). A 20x (0.70 NA) objective and additional 4x optical zoom were used for particle tracking, capturing an average of 15 ± 5 trackable particles in 30 second videos at 26 frames per second. For the long-term timelapse imaging, ten organoids were imaged over 72 hours at 5 min intervals. For particle tracking microrheology in native gastric mucus (pH 6.5±0.5) or PGM (15 mg/mL (14, 61)), mucus and 1 µm Fluoresbrite polystyrene microspheres were pipetted into 0.12 mm spacers (SecureSeal™, Grace Bio-Labs, Bend, OR) that were mounted on glass slides and then sealed with glass cover slips. A Keyence BZ-X810 microscope with a 60x oil-immersion objective (1.4 NA) was used for imaging. For each sample, we recorded six 30 second videos at a frame rate of 29 frames per second and a frame size of 640 x 480 pixels at room temperature.

Videos were analyzed in MATLAB v7.9.0 using the “polyparticletracker” particle tracking routine, which finds the center of intensity for each particle using a polynomial Gaussian fit (72). Drift was corrected, if necessary, using a 3D Drift Correction plugin for ImageJ (73) before analyzing the videos. Parameters within the particle tracking software were adjusted to only track singlets of particles. Using the trajectory data for each particle, the time-dependent mean square displacement was calculated:

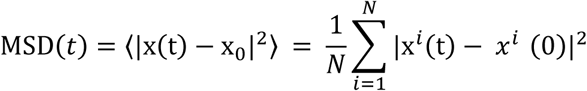

in which N = total number of particles to be averaged, x*^i^* (0) = initial particle reference position, and x*^i^* (*t*) = particle position at time *t*.

An exponential model was fit to each tracked MSD profile and power constant α was recorded:

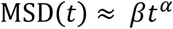

in which α = exponential slope of the MSD profile. In the instance ɑ ≈ 1.00, linear regression was conducted to determine the average slope of MSD(*t*) and subsequently used to calculate the diffusion coefficient using the generalized Stokes-Einstein equation (GSE) (74, 75). G^∼^(*s*) was calculated by taking the unilateral Laplace transform of MSD(*t*) for each particle using the GSE. Allowing *s* = i*ω*, we obtained frequency-dependent storage (elastic) (G’) and loss (viscous) (G’’) moduli (74).

### Monolayer Culture

Transwell permeable supports with PET membranes (surface area: 0.33 cm^2^; pore diameter: 0.4 µm; Corning, 3470) were coated with 15 µg/cm^2^ rat tail collagen I (Corning, 354236) for 1 hour at room temperature(20, 21). After aspirating off the remaining collagen, the basolateral chambers were then filled with 600 µL L-WRN culture medium and incubated for at least one hour at 37°C 5% CO_2_. Human gastric organoids (cultured as described above) were expanded, harvested, and trypsinized for 10 minutes in a 37°C water bath (21). The resulting organoid fragments were mechanically dissociated as previously described, and pipetted through a 70 µm cell strainer to eliminate cell clusters (68). The filtrate was centrifuged at 400 x *g* for 5 minutes and the resulting cell pellet was resuspended in L-WRN. A minimum of 2.0 × 10^5^ cells were seeded onto each insert, agitated gently for 5 minutes, and incubated undisturbed at 37°C 5% CO^2^ for at least 24 hours. Every other day, media was replenished and transepithelial electrical resistance (TEER) was measured using an EndOhm Chamber attachment to an EVOM 2 epithelial voltohmeter (World Precision Instruments). Once a transepithelial electrical resistance (TEER) of at least 200 Ω*cm^2^ was reached, apical medium was removed to begin culture at the air-liquid interface (ALI)–a process termed “airlifting”. Cells were then left undisturbed until there was visible production of at least 50 µL apical mucus for collection. Our gastric epithelial cells typically form confluent monolayers within one week, and mucus production was observed within one week of airlifting.

### Harvesting and Storage of Bioengineered Gastric Mucus

Bioengineered mucus (BGM) was harvested off the apical epithelium at either biweekly or weekly intervals using either forceps or a wide-bore pipette and transferred to a microcentrifuge tube. The wet weight of the mucus harvested from each well was recorded. Samples were then either used immediately for experiments, frozen at −20°C, or frozen at −80°C for long-term storage. For most experiments, BGM and native mucus (NM) samples were used fresh to preserve their natural state as much as possible (76). Samples for which dry weight was measured were lyophilized, as described below. Porcine gastric mucin (PGM), prepared as previously described (77), was lyophilized and rehydrated at a concentration of 10-15 mg/mL for experiments (78).

### Mucus Lyophilization

BGM samples were first flash-frozen in liquid nitrogen and stored on dry ice for a minimum of 30 min prior to lyophilization for at least 48 h with a FreeZone^TM^ benchtop freeze dryer (Labconco^TM^ 700202000). The dry weight of each sample was recorded for quantification of dry mass and water content. Lyophilized BGM samples were only used for the determination of dry mass.

### MUC5AC ELISA

For quantification of MUC5AC protein concentration in the NM and BGM samples, a human extracellular MUC5AC ELISA kit (Abcam, ab303761) was used following the manufacturer’s protocol. As not every monolayer yielded sufficient mucus at each collection time point, samples were pooled so that each sample analyzed via ELISA represented the yield from three Transwells. BGM samples included both lyophilized and non-lyophilized samples that had been stored at −20°C. Organoid culture media saved from “blank” Transwells and stored under the same conditions was used as negative control.

### Size Exclusion Chromatography

Samples for size exclusion chromatography (SEC) were prepared by dissolving 50 μL of mucin in 150 μL of buffer, followed by filtration through a 0.45 μm syringe filter. The buffer comprised 0.001 M Na_2_HPO_4_, 0.001 M EDTA, 0.1% (w/v) NaN_3_, and 0.03 M NaCl. BGM samples were diluted 1:1. A volume of 20 μL from each sample was then loaded onto a YMC Diol-300 SEC column at a flow rate of 1 mL/min. Molecular weight determination and concentration measurements were conducted using multi-angle light scattering (MALS) and refractive index (RI) detection, respectively, through in-line Wyatt Detectors. All data sets were analyzed using Astra 8 software and normalized against commercial PEG and PEO standards from Wyatt.

### Proteomics

Proteomic analysis was performed at the GlycoMIP core facility at Virginia Tech using native human gastric mucus, BGM, and porcine gastric mucin. Cysteine disulfide bonds were reduced using 4.5 mM dithiothreitol (DTT) and incubation at 37°C for one hour. Free sulfhydryl groups were alkylated with 10 mM iodoacetamide (IAA) at room temperature for 30 min in the dark, then unreacted IAA was quenched with 10 mM DTT. Protein was precipitated by the addition of *o*-phosphoric acid to 1.2% (v/v) and 1 mL methanol followed by overnight incubation at −80°C. Precipitated protein was loaded onto a micro S-Trap (Protifi) by centrifugation at room temperature for 1 minute at 1,000 x *g* and washed extensively with methanol. Samples were digested using 2 µg. Pierce Trypsin Protease, MS grade (ThermoFisher Scientific) in 50 mM triethylammonium bicarbonate (pH 8.5) at 37°C overnight. Peptides were recovered by sequential washings of the S-Trap (25 µL solvent A, 25 µL 50:50 solvent A: solvent B, 25 µL solvent B). Excess acetonitrile was removed by vacuum centrifugation and peptide concentrations were determined using a nano-UV/Vis spectrometer (DeNovix) to measure the absorbance at 215 nm. (Solvents A and B are the same as solvents A and B used for the chromatography.) The processed samples were analyzed by LC/MS on a Bruker MALDI-2 Mass Spectrometer (timsTOF Pro/flex) equipped with a Vanquish Neo UPLC unit. The gradient used LCMS-grade water with 0.01% formic acid as solvent A and 80% acetonitrile with 0.01% formic acid as solvent B. The column used was a μPACTM HPLC column (ThermoFisher Scientific, catalog number COL-NANO200G1B), and the flow rate was 8 μL/min. The run was 95 min long. The starting percentage of solvent B was 2%. The final percentage of solvent B was 98%.

Proteins identified as “reverse sequences” and those flagged as “common contaminant proteins”, such as porcine albumin, were excluded from the data set. Protein isoforms were treated as separate entries. Proteins were categorized based on the human protein atlas (79) as (i) mucins; (ii) gastric proteins, i.e., secreted proteins from the GI tract or proteins known to have a specific function in the stomach; (iii) cellular proteins, i.e., cytoplasmic, nuclear, or membrane proteins with no specific known function in the stomach; (iv) contaminants, i.e., proteins derived from blood/serum or extracellular matrix; or (v) unknown proteins.

### CryoFE-SEM

Cryo-Field Emission Scanning Electron Microscopy (cryoFE-SEM) was performed at the Imaging and Chemical Analysis Laboratory (ICAL) at Montana State University. NM from three tissue donors and BGM samples from three different organoid lines were analyzed. Approximately 50 µL of mucus (or BGM) were sandwiched from atop the ALI culture directly between two gold-coated (Emitech K575X) silicon wafers, flash-frozen in liquid nitrogen, and manually fractured very quickly along scribed lines on the wafers using a flathead screwdriver before imaging on a Zeiss SUPRA 55VP Field Emission Scanning Electron Microscope. Images were acquired at −140°C with secondary electrons, both in high vacuum and variable pressure modes, at accelerating voltages of 0.8-1 kV and a 10 µm aperture.

Image analysis was performed in FIJI using a Segment Anything Model (SAMJ; **Supp. Fig. 3**)(41). Images were first converted to grayscale and automatically enhanced for contrast and brightness. Fully automated segmentation was not feasible due structural heterogeneity of the samples due to debris and other artefacts, particularly in the native mucus samples. These factors resulted in frequent false positives and fragmented boundaries when using automated threshold-based methods. Instead, SAMJ enabled a semi-automated workflow consisting of user-imposed prompts, generation of binary masks, and optional model training via re-selection of incorrectly recognized ROIs. The EfficientViTSAM image was selected for its cross-platform compatibility, particularly important for low-GPU devices. Structures of interest (intact honeycombs or pores) were first manually localized using bounding boxes drawn with the rectangle tool, after which SAMJ automatically delineated boundaries. Such an approach reduces operator bias compared to manual tracing while avoiding errors inherent to threshold-only segmentation. Identified ROIs were measured and exported as binary masks. To distinguish honeycomb cavities from pores, ROI measurements were gated based on FIJI circularity and solidity descriptors, applying thresholds derived from receiver operating characteristic (ROC) analysis of manually annotated ground-truth structures (**Supp. Fig. 2A**). In FIJI, a circularity is 4π(area/perimeter^2^) (with 1.0 indicating a perfect circle), while solidity is the ratio of area to convex hull area (with lower values indicating more concave or irregular boundaries). For both honeycomb and pore area measurements, outliers were identified and excluded using the ROUT method (Q = 1%). Measurements were compared between native and bioengineered mucus sample types using unpaired t-tests.

### Bulk Rheometry

For rheological measurements, all mucus samples were equilibrated to room temperature. Storage (elastic, G’) and loss (viscous, G’’) moduli were measured via the following series of rheometric tests: (I) flow sweeps to obtain viscosity as a function of shear rate, (II) amplitude sweeps to determine the linear viscoelastic region in which subsequent measurements are performed, and (III) frequency sweeps to determine the viscoelastic moduli(80, 81). All tests were performed using an AR-G2 rheometer with a 20 mm parallel plate geometry (TA Instruments)(82, 83). An initial flow sweep was performed to obtain steady shear viscosity as a function of shear stress. Next, an amplitude sweep was performed in a range starting just before the point where the viscosity began to drop off in the flow sweep. The amplitude sweeps were performed at a constant frequency of 1 Hz with oscillatory shear stress of increasing amplitude to determine the linear viscoelastic region (LVR) for each sample. The LVR was determined by identifying the region in which the G’ and G’’ plateau because they are undisturbed by the oscillatory strain(84). A strain amplitude of 1% within this linear regime was then used for subsequent frequency sweeps (0.1 – 2.5 rad/s) of the samples. Samples were allowed to recover for 10 minutes in between tests(84).

### Consideration of biological variables, rigor and reproducibility, and statistical analysis

Tissue samples were obtained from donors of any sex or ethnicity and within an age range of 18-70. Each experiment was repeated three or more times with different organoid lines or native mucus samples. All data were analyzed using GraphPad Prism version 10.2.3 (San Diego, CA, USA). Data are presented as the mean ± SD. Student’s *t* tests and one-way ANOVA with Tukey’s or Dunnett’s multiple comparisons tests for normally distributed data or the Kruskal-Wallis test with Dunn’s multiple comparison test for data without Gaussian distribution were used to assess statistical significance. *P* ≤ 0.05 was considered to indicate a statistically significant difference. The ROUT coefficient method with a Q value of 1% was used to remove outliers.

## Supporting information

Supplementary data 1 proteomics

## Ethics Statement

Human gastric tissue samples were obtained with informed consent and Institutional Review Board (IRB) approval from patients undergoing endoscopy and biopsy at the Bozeman Health Deaconess Hospital (protocol 2023-48-FCR). Alternatively, de-identified whole stomachs collected post-mortem from transplant donors or surgical discard material from sleeve gastrectomy were provided by the National Disease Research Interchange. Those samples were considered exempt from IRB review (protocol DB062615-EX).

## Author Contributions

KNL: conceptualization, formal analysis, methodology, investigation, visualization, funding acquisition, writing: original draft, writing: review, and editing; BS: conceptualization methodology, investigation, visualization, writing: original draft; CD: investigation, formal analysis, visualization, writing: original draft; CV: investigation, formal analysis; GJ: investigation; TAS: investigation; AB: investigation; LD: methodology, investigation; RFH: methodology, investigation, formal analysis; WL: methodology, BD: methodology, investigation; JB: methodology, resources; SGM: methodology, supervision, funding acquisition; JNW: conceptualization, methodology, supervision, funding acquisition, writing: original draft; RB: conceptualization, methodology, resources, supervision, funding acquisition; DB: conceptualization, methodology, formal analysis, visualization, supervision, project administration, funding acquisition, writing: original draft, writing: review, and editing. All authors reviewed and approved the final version of the manuscript.

## Data Availability Statement

The datasets generated during the current study are included in this submission or are available from the corresponding author on reasonable request. Proteomics datasets will be made available through the PRIDE data repository.

## Acknowledgements

This project was supported by the National Institutes of Health, awards R01GM131408 (D.B., J.N.W., R.B.), U01EB029242 (D.B. and J.N.W.), and UL1 TR002319 (Research Training Fellowship to K.N.L.), and by the National Science Foundation, awards DMS-1813654 (S.G.M.) and CBET-2112085 (S.G.M.). Access to CryoFE-SEM was provided by a user grant to D.B. from the Montana Nanotechnology Facility (MONT), which is supported by the National Science Foundation (Grant# ECCS-2025391). This work was supported by GlycoMIP, a National Science Foundation Materials Innovation Platform funded through Cooperative Agreement DMR-1933525. The authors would like to thank Dr. Clover Su for helpful discussions regarding experimental methodologies, and Dr. Bradley S. Turner for providing purified porcine gastric mucin and for helpful discussions.

## Supplementary Material

### Supplementary Figure Legends

**Supplementary Figure 1.**
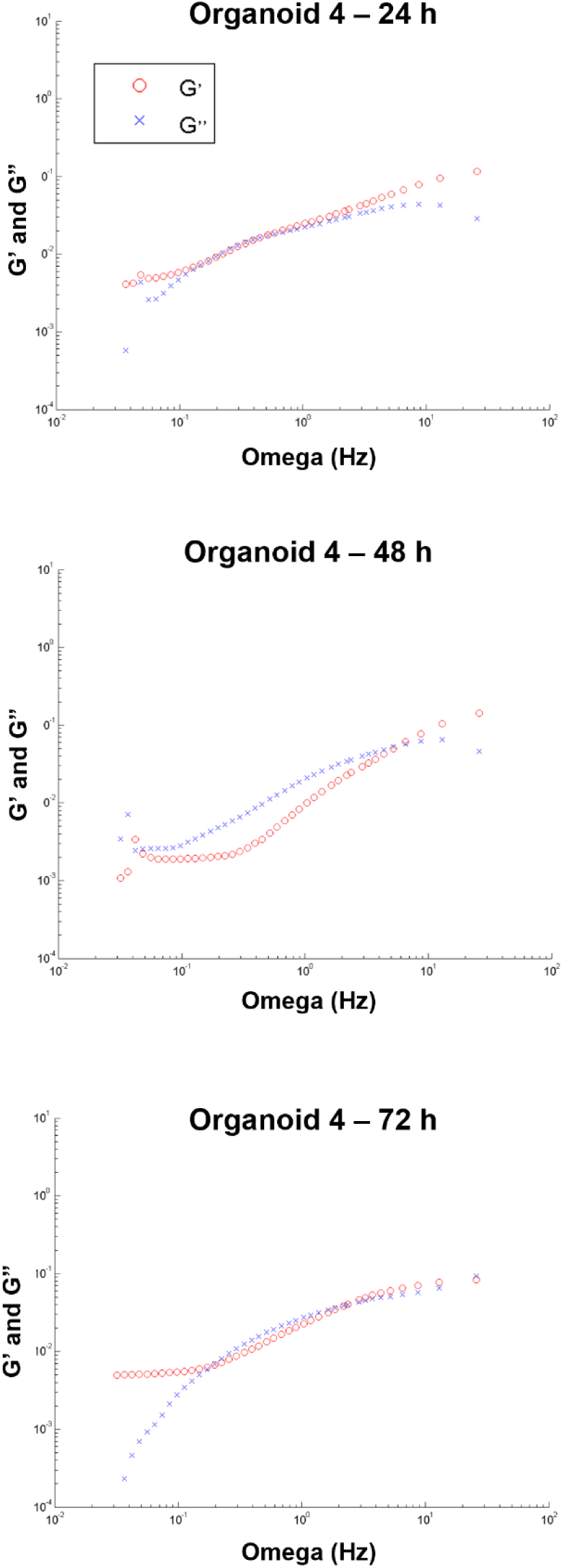
Angular frequency-dependent elastic (G’) and viscous (G”) moduli for mucus in the representative organoid shown in Fig. 2. The organoid was injected with 1 µm beads and was analyzed at 24 h, 48 and 72 h post injection by particle tracking microrheology. Elastic moduli (G’) represented by red dots; viscous moduli (G”) represented by blue crosses.

**Supplementary Figure 2.**
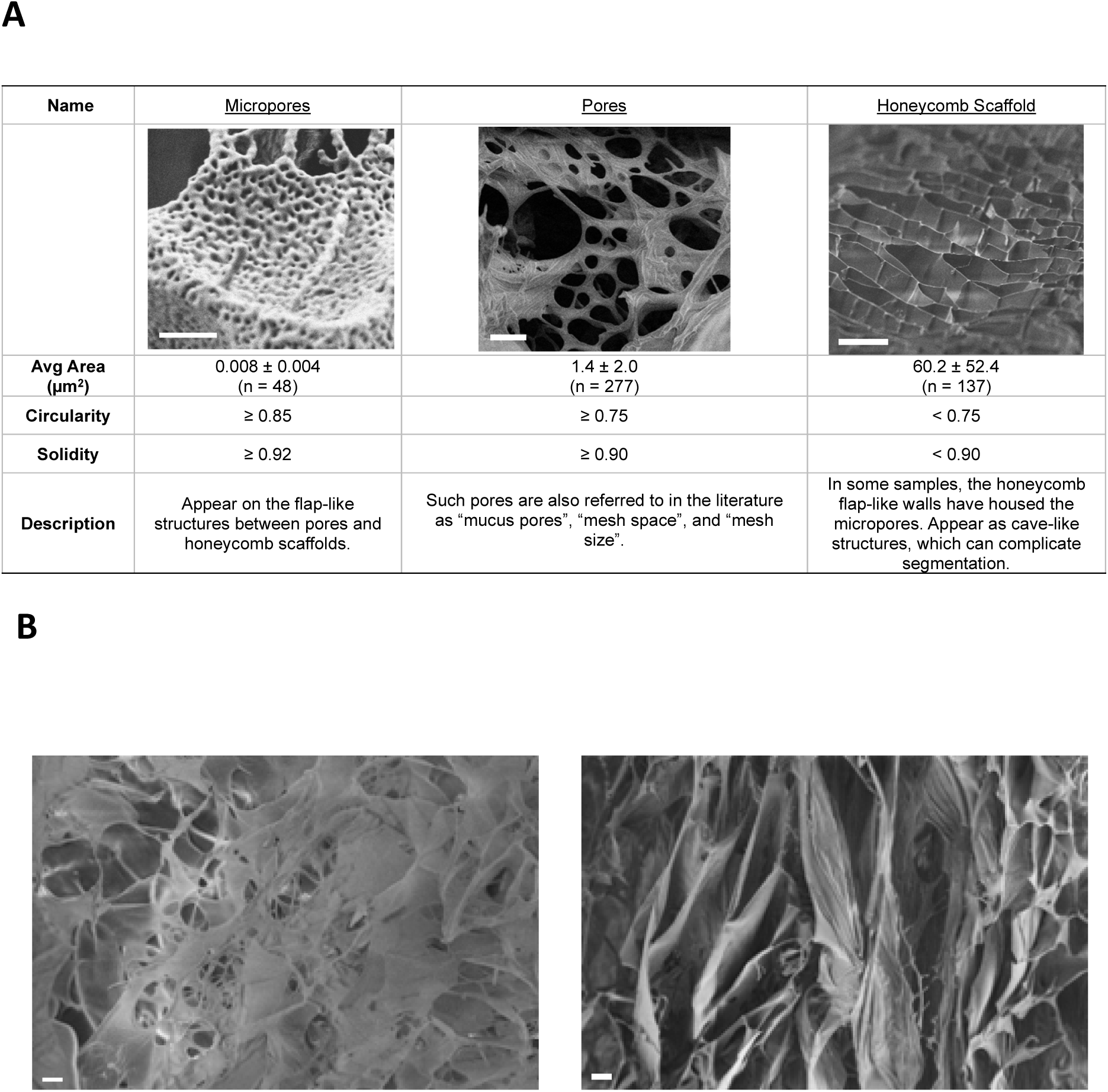
(**A**) Categories and criteria for different structures identified in gastric mucus samples by CryoFE-SEM. Micropores, observed in bioengineered (n = 2) and native (n = 2) mucus samples (scale bar = 1 µm). Pores, observed in bioengineered (n=3) and native (n = 1) mucus samples (scale bar = 20 µm). Honeycomb scaffold, observed in bioengineered (n = 2) and native (n = 2) mucus samples (scale bar = 20 µm). (**B**) Controls for CryoFE-SEM. (left panel) L-WRN culture media. (right panel) L-WRN culture media collected from a blank, collagen-coated Transwell.

**Supplementary Figure 3.**
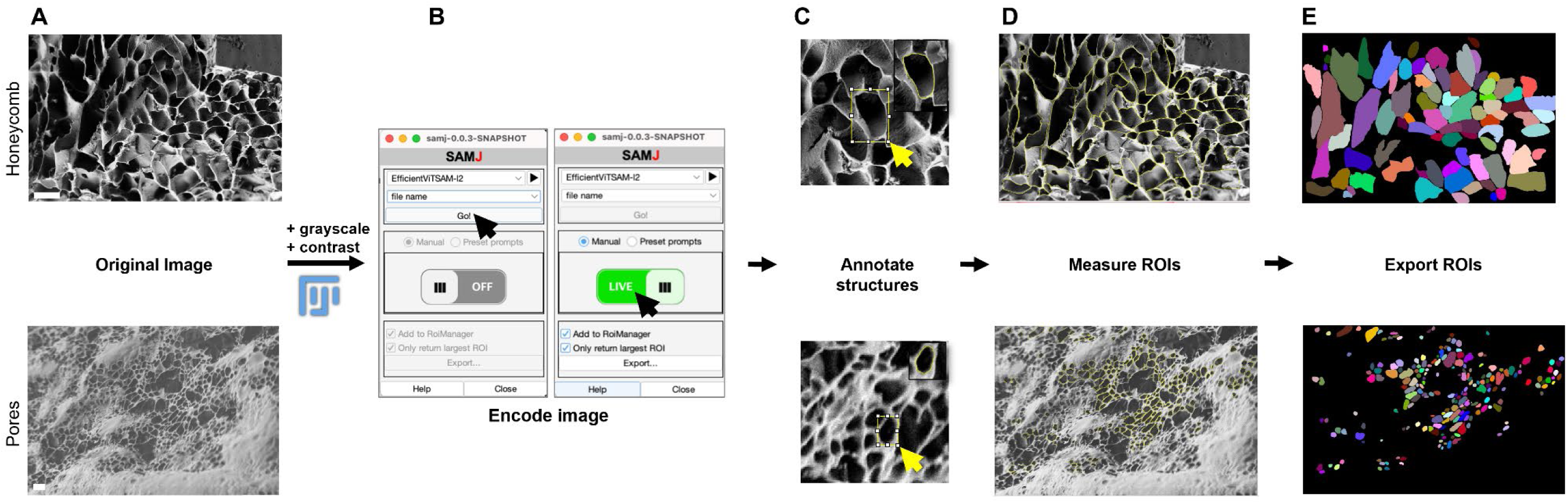
Image analysis pipeline for measurement of honeycomb and pore sizes using Segment Anything Model (SAMJ)(41). (**A**) Original images are uploaded to FIJI in TIF-format, converted to grayscale, and auto-corrected for brightness and contrast. Scale bars correspond to 10 µm for honeycomb (top) and 2 µm for pores (bottom). (**B**) SAMJ plugin is opened and image encoder is selected (in this case: EfficientViTSAM-i2, which is compatible with most computing setups; left). Annotation mode is set to “Live” (right). (**C**) The rectangle selection tool is used to manually create a bounding box around the structure of interest–a prompt that is subsequently processed by the model. (**D**) Almost instantaneous generation of binary masks corresponding to each prompt (bounding box). Inset shows the auto-generated mask based on the manually defined bounding box. (**D**) Overlay of all measured ROI outlines. (**E**) Exported binary masks.

**Supplementary Table 1:**
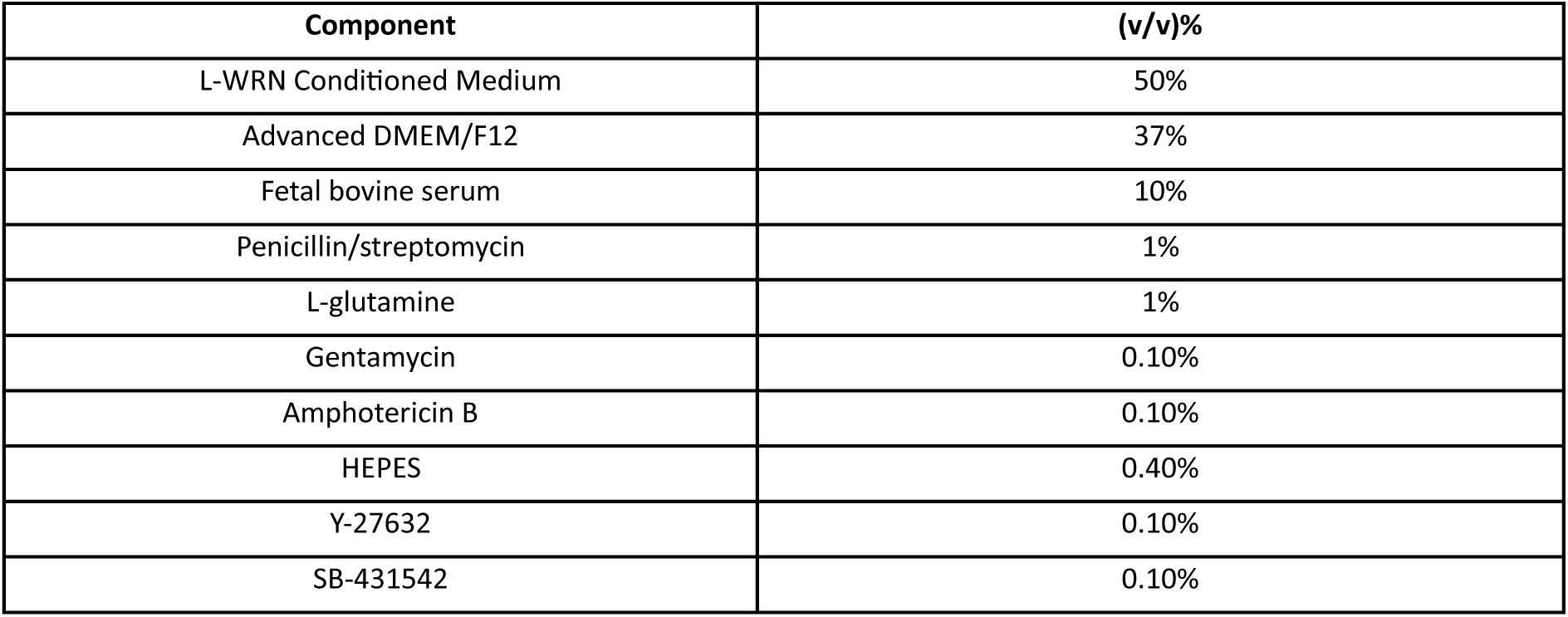
Human gastric organoid expansion media components.

**Supplementary data 1: Complete proteomics data set and lists of top 35 proteins**

Excel spreadsheet with (a) complete proteomics dataset organized by sample and (b) list of top 35 highly expressed proteins present in all samples in each category (BGM, NM, PGM).

